# Longitudinally Mapping Childhood Socioeconomic Status Associations with Cortical and Subcortical Morphology

**DOI:** 10.1101/352187

**Authors:** Cassidy L. McDermott, Jakob Seidlitz, Ajay Nadig, Siyuan Liu, Liv S. Clasen, Jonathan D. Blumenthal, Paul Kirkpatrick Reardon, François Lalonde, Raihaan Patel, Mallar M. Chakravarty, Jason P. Lerch, Armin Raznahan

## Abstract

Childhood socioeconomic status (SES) impacts cognitive development and mental health, but its association with structural brain development is not yet well-characterized. Here, we analyzed 1243 longitudinally-acquired structural MRI scans from 623 youth to investigate the relation between SES and cortical and subcortical morphology between ages 5 and 25 years. We found positive associations between SES and total volumes of the brain, cortical sheet, and four separate subcortical structures. These associations were developmentally fixed rather than age-dependent. Surface-based shape analysis revealed that higher SES is associated with areal expansion of (i) lateral prefrontal, anterior cingulate, lateral temporal, and superior parietal cortices and (ii) ventrolateral thalamic, and medial amygdalo-hippocampal sub-regions. Meta-analyses of functional imaging data indicate that cortical correlates of SES are centered on brain systems subserving sensorimotor functions, language, memory, and emotional processing. We further show that anatomical variation within a subset of these cortical regions partially mediates the positive association between SES and IQ. Finally, we identify neuroanatomical correlates of SES that exist above and beyond accompanying variation in IQ. Our findings clarify the spatiotemporal patterning of SES-related neuroanatomical variation and inform ongoing efforts to dissect the causal pathways underpinning observed associations between childhood SES and regional brain anatomy.

## Introduction

Early brain development occurs within the context of each child’s experiences and environment, which both vary significantly as a function of socioeconomic status (SES). SES is a multidimensional construct that reflects an individual or family’s access to economic and social resources. Childhood SES is typically measured by factors including family income, parental education, and parental occupation (Hollingshead 1957; McLoyd 1998). Childhood SES, across a wide range of levels, has been associated with disparate outcomes in mental health, cognitive development, and academic achievement (Brooks-Gunn and Duncan 1997; Sirin 2005; Noble et al. 2007; Reiss 2013). These associations are thought to arise through diverse causal pathways, including (i) direct SES-linked environmental effects on cognitive outcomes (Ritsher et al. 2001; Kendler et al. 2015), (ii) the capacity of mental health difficulties or low cognitive performance to negatively impact SES (Tiikkaja et al. 2016), and (iii) the existence of factors that simultaneously increase risk for lowered SES and cognitive difficulties (Trzaskowski et al. 2014; Hill et al. 2016).

The robust epidemiological data connecting childhood SES to behavioral and cognitive development have motivated a series of recent neuroimaging studies aimed at mapping SES effects on both global and regional brain anatomy. Higher SES has been associated with greater total grey matter volume, and less consistently with white matter volume (Luby et al. 2013; Hair et al. 2015; Gianaros et al. 2017), as well as greater volume in certain cortical and subcortical sub-regions of a priori interest, including in particular the prefrontal cortex and the hippocampus (Jednoróg et al. 2012; Noble et al. 2012; Hanson et al. 2013; Luby et al. 2013; Hair et al. 2015; Holz et al. 2015). Furthermore, a recent landmark study by Noble and colleagues (Noble et al. 2015) mapped spatial associations between socioeconomic factors and cortical surface area using cross-sectional neuroimaging data. These pioneering studies suggest that observed associations between childhood SES and neurodevelopmental outcomes may be centered on specific brain systems, and raise as yet unresolved questions regarding when SES-neuroanatomy associations are established, whether these associations are predominantly cortical or subcortical in nature, and how the observed correlations between SES and brain anatomy might be related to accompanying variation in cognitive ability (Noble et al. 2007; for review, see: Brito and Noble 2014). Addressing these open questions would represent an important step forward in detection of biological pathways underlying the capacity of childhood SES to predict later health.

Here, we seek to build on our current understanding of the relation between SES and brain development in five key directions. First, to determine if SES associations with brain anatomy are dynamic or fixed across development (Giedd et al. 1999), we model SES-brain associations within a large, longitudinal sample of over 1200 structural neuroimaging scans from over 600 healthy individuals spanning 5 to 25 years of age. Second, within the cortical sheet, we separately model SES associations with regional cortical thickness (CT) and surface area (SA) – two biologically distinct cortical phenotypes which together determine cortical volume (Raznahan et al. 2011). Findings remain sparse and mixed regarding the relative strength and spatial distribution of SES relations with SA and CT (Lawson et al. 2013; Noble et al. 2015). Third, we extend our analyses beyond the cortical sheet to systematically assess the relation between SES and the anatomy of five major non-cortical structures (henceforth referred to as “subcortical” structures) using multi-atlas methods that provide both estimates of bulk volume and spatially fine-grained measures of each structure’s shape. Simultaneous examination of SES associations with regional anatomy of the cortex and subcortex is critical given evidence that these brain systems are known to function (Redgrave et al. 2010), develop (Raznahan et al. 2014) and connect (Draganski et al. 2008) with each other in a topographically organized manner. Fourth, we formally characterize the functional associations of those cortical regions which are anatomically correlated with SES using the Neurosynth platform for online meta-analysis of neuroimaging data (Yarkoni et al. 2011). Finally, given known positive associations between SES and cognitive performance (Brooks-Gunn and Duncan 1997; Kendler et al. 2015) and the potential of brain anatomy to vary as a function of cognitive ability (Walhovd et al. 2016), we probe the complex associations between SES, neuroanatomy, and cognition through two methods that have been used separately in previous literature, namely: (i) including IQ as a covariate in models of SES effects on brain anatomy (Noble et al. 2012; Lawson et al. 2013) and (ii) for all structural phenotypes that show a relation with SES, assessing whether these anatomical disparities mediate the relation between SES and IQ (Hair et al. 2015; Noble et al. 2015).

## Materials and Methods

### Participants

This study includes a longitudinal sample of 1243 structural magnetic resonance imaging (sMRI) brain scans from 623 healthy children and adolescents between 5 and 25 years old (Table 1). Participants were recruited through local advertisement for a study of typical brain development conducted at the National Institute of Mental Health Intramural Research Program between 1990 and 2010. Participants were screened and excluded on the basis of a history of mental health treatment, use of psychiatric medication, enrollment in special services at school, or diagnosis of any medical condition known to affect the nervous system. The research protocol was approved by the institutional review board at the National Institute of Mental Health, and written informed consent or assent was obtained from all children who participated in the study, as well as consent from their parents if the child was under the age of 18.

**Table 1.** Participant Characteristics

| Characteristic | Data |
| --- | --- |
| No. of individuals | 623 |
| No. of scans | 1243 |
| Sex, no. |  |
| Female | 299 |
| Male | 324 |
| Age at first scan, years |  |
| Mean (SD) | 12.0 (4.0) |
| Range | 5.2 – 25.4 |
| Handedness, no. |  |
| Left | 35 |
| Right | 537 |
| Mixed | 51 |
| Race, no. |  |
| White | 534 |
| Black | 40 |
| Asian | 14 |
| Hispanic | 20 |
| Other | 15 |
| IQ |  |
| Mean (SD) | 114 (12.4) |
| Range | 78 – 150 |
| Hollingshead SES |  |
| Mean (SD) | 41.2 (18.3) |
| Range | 95 – 20 |
| No. of scans per individual, no. |  |
| 1 scan | 279 |
| 2 scans | 168 |
| ≥ 3 scans | 176 |

### Socioeconomic Status

Childhood socioeconomic status (SES) was quantified using the Amherst modification of the Hollingshead two-factor index (Hollingshead 1957; Watt 1976). Parental education and occupation were each coded on a 7-point scale, and these two values were used to derive and record a single SES score that was used for analyses (for occupation and education categories and scoring, see Tables S1 and S2). When education and occupation were reported for two parents, the highest SES was used. In the resulting index, children from the most advantaged families receive the lowest Hollingshead score of 20, while children from the most disadvantaged families receive the highest Hollingshead score of 134. For ease of interpretation, we refer to SES variation using conventional directionality, such that “higher SES” refers to a lower Hollingshead score. Accordingly, reported positive associations with SES in the manuscript indicate variables that increase in value with greater SES, and are thus negatively correlated with Hollingshead score. In graphical representation of such associations, x-axis scales for SES are reversed, such that right-most values index the highest SES levels (i.e., lowest Hollingshead scores).

### General Cognitive Ability

Full-Scale IQ was estimated for each child in the sample using an age-appropriate Wechsler scale. The majority (*N* = 562) of children received the Wechsler Abbreviated Scale of Intelligence (Wechsler 1999), and other scales used include the WAIS-R, WISC-R and WISC-III, and the WPPSI and WPPSI-III. For children who were tested multiple times, the most recent, complete IQ measurement was used.

### Image acquisition and processing

All sMRI scans were T1-weighted images collected on the same 1.5 T General Electric SIGNA scanner with contiguous 1.5 mm axial slices using a 3D spoiled-gradient recalled-echo sequence (echo time = 5ms; repetition time = 24ms; flip angle = 45°; acquisition matrix = 256 × 192; number of excitations = 1; field of view = 24 cm). All scans were visually inspected to ensure limited motion artifact.

Native sMRI scans were submitted to the CIVET 1.1.10 pipeline for automated morphometric analysis (Ad-Dab’bagh et al. 2006). The CIVET pipeline uses a validated neural net approach to voxel classification to calculate grey matter and white matter volume estimates (Zijdenbos et al. 2002; Cocosco et al. 2003) after initial correction of images for radiofrequency intensity non-uniformities (Collins et al. 1994; Sled et al. 1998). Following tissue classification, each image was fitted with two deformable mesh models to identify the inner and outer surface of cortical grey matter, and these surfaces were used to calculate cortical thickness and surface area at 40,962 vertices on each cortical hemisphere as previously described (MacDonald et al. 2000; Raznahan et al. 2013).

Subcortical segmentation and surface extraction were completed automatically using the MAGeT Brain algorithm (Chakravarty et al. 2013; Raznahan et al. 2014). Scans were first registered to the ICBM 152 template and corrected for radiofrequency intensity non-uniformities (Collins et al. 1994; Sled et al. 1998). For the striatum, thalamus, and pallidum, the segmentation atlas was created using a 3D reconstruction of serial histological data, warped to an MRI-based template (Chakravarty et al. 2006). The MAGeT pipeline then customized this atlas to 21 randomly selected subjects within the sample. All 1243 scans were then warped to this set of templates, providing a set of 21 candidate subcortical segmentations for each scan. For the hippocampus and amygdala, five reference atlases were generated from high resolution and contrast T1 and T2 weighted images from three males and two females using a 3T scanner with final super-sampled isotropic voxel dimensions of 0.3 mm (Wood 2011). The MAGeT pipeline again created automated segmentation atlases for 21 randomly selected subjects, resulting in 105 possible segmentations (5 atlases × 21 templates) for each of the 1243 scans in our dataset. Each scan was labeled using the 21 striatum, thalamus and pallidum segmentations and the 105 hippocampus and amygdala segmentations, and the final segmentation was decided upon using a label voting procedure, such that the label occurring most frequently at each voxel was retained. These procedures provided estimates of total bilateral volume for the hippocampus, amygdala, thalamus, striatum and pallidum in each scan. All scans used for analysis passed visual quality control of these final subcortical segmentations to exclude visible segmentation errors of the five subcortical structures under study.

Surface-based representations of all five subcortical structures were then estimated on their respective atlases using a marching cubes algorithm (Lerch et al. 2008). Next, the nonlinear portions of the 21 transformations mapping each subject to the 21 input templates were concatenated and averaged across the template library to limit noise and increase precision and accuracy. These surface-based representations were warped to fit each template and each surface was warped to match each subject. This procedure yields 21 possible surface representations per subject for the striatum, thalamus and pallidum, and 105 possible surface representations for the hippocampus and amygdala, which were merged by estimating the median coordinate representation at each location. Next, a third of the surface area of each triangle forming the surface representation was assigned to each vertex within the triangle. The surface area at each vertex is the sum of all such assignments from all connected triangles. Finally, surface area values were blurred with a surface-based diffusion-smoothing kernel (5mm for the amygdala, hippocampus, striatum and thalamus, and 3mm for the pallidum). This processing stream generated surface area values for a total of 26,401 vertices across the five subcortical structures in each scan.

### Statistical analyses

The effect of SES on each anatomical metric of interest was modeled using a linear mixed effects model with sex and centered age as fixed effects covariates and each individual’s ID (to account for multiple longitudinal scans per individual) and family ID (to account for the presence of dizygotic twin pairs and siblings in the sample) as nested random effects. The decision to present core results with SES, sex and age as main effects was made after first ruling out the presence of extensive interactions between these variables in predicting structure or vertex-level anatomical variation effect on each anatomical variable. The few cases where interactions were found will be noted in the text. Otherwise, anatomy for the *i*th family’s *j*th individual’s *k*th time point was modeled as follows:

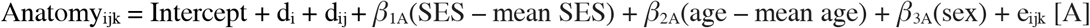

Dependent volumetric variables of interest included total brain volume (TBV – the sum of intracranial grey and white matter volume), total intracranial grey matter volume (GMV) and total intracranial white matter volume (WMV), the total bilateral volume of the cortex (CV), and total bilateral volume of each subcortical structure (i.e., “bulk” hippocampal, amygdalar, thalamic, striatal, and pallidal volumes). To aid comparison of the associations between SES and these diverse volumetric indices, we also estimated the standardized effect for each SES-volume association by re-running model [A] above after centering and scaling all variables so that the resulting β_1_ coefficient would index the standard deviation shift in volume with one standard deviation increase in SES (i.e., a decrease of approximately 18 Hollingshead points).

Vertex-level anatomical variables of interest included cortical thickness (CT) and surface area (SA) at each of 80,962 cortical vertices and SA at each of 26,401 subcortical vertices (hippocampus, 1152 left/1215 right; amygdala, 1473 left/1405 right; thalamus, 3016 left/3108 right; striatum, 6450 left/6178 right; pallidum, 1266 left/1138 right). Vertex-specific β_1_ coefficients were visualized on the corresponding cortical or subcortical surface after applying a false discovery rate (FDR) correction for multiple comparisons. FDR corrections were calculated separately across the left and right cortical hemispheres and the left and right subcortical structures with *q* (the expected proportion of false rejections of the null hypothesis) set to 0.05.

Finally, we probed the relation between SES, anatomical metrics, and cognition in two ways. First, for all structures that showed a significant main effect of SES in model [A], we tested the robustness of this effect to the inclusion of Full-Scale IQ as a covariate in the linear mixed effects models. Separate main effects of SES and IQ were included after first ruling out the presence of a significant SES by IQ interactive effect. For these analyses, anatomy for the *i*th family’s *j*th individual’s *k*th time point was modeled as follows:

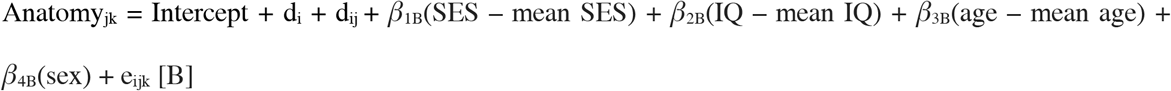

Linear age terms were used in models [A] and [B] after verifying that this simple parameterization of age yielded identical SES findings to models run with age parameterized as a non-linear spline (which could allow for non-linear age effects). Additionally, models [A] and [B] were re-run for each anatomical metric within the subset of participants (n = 534) who self-endorsed the federal race category of “white,” and all main effects held.

Finally, we conducted mediation analyses to investigate the extent to which the global and vertex-wise structural associations with SES mediate the association between SES and IQ. For the mediation tests with a single global mediator, we used the Mediation package in R (Tingley et al. 2014), with SES as the independent variable (treatment) and Full-Scale IQ as the dependent variable. The *mediate()* function estimates the average causal mediation effect (ACME) and the average direct effect (ADE), which together sum to the total effect of the treatment (i.e., SES) on the outcome (i.e., IQ). The proportion mediated, which we report, represents the size of the ACME relative to the total effect. Within the map of cortical vertices that showed a significant association with SES, we tested the mediating role of each vertex using the MultiMed package in R (Boca and Sampson 2014), which implements a permutation approach with joint correction to test multiple mediators simultaneously.

### Interpretation of Anatomical Results

To systematically investigate the functional implications of SES effects, we submitted the MNI coordinates of peak SES effects on surface area and cortical thickness to Neurosynth (Yarkoni et al. 2011). Neurosynth is an online platform that extracts and synthesizes brain activation patterns and psychological terms across >11,000 functional neuroimaging publications. It can be used to generate correlations between the meta-analytic co-activation map for a given point of interest in the brain and each of the terms in the Neurosynth database. Using Neurosynth, we identified cognitive and psychological terms that frequently co-occur in the literature with functional activations similar to the observed pattern of SES effects on cortical morphology.

## Results

### Participant Characteristics

Participant characteristics are detailed in Table 1. Hollingshead SES in our sample ranged from a high of 20 to a low of 95 after 3 scans from individuals with exceptionally low SES scores (*z* = 4) were removed. The complete Hollingshead scale ranges from 20 to 134, and approximately one-quarter of our 623 participants received the highest SES score of 20. Thus our sample’s distribution of Hollingshead scale scores does not include the highest values possible (which would reflect the most severe levels of socioeconomic disadvantage), and the concentration of individuals with the lowest score possible (20) suggests an enrichment relative to the general population for children with parents who received a graduate professional degree and employment in the highest occupation category (see Tables S1 and S2). Throughout the text below, SES is referred to using conventional directionality, such that “higher SES” refers to a lower Hollingshead score, and reported positive associations with SES are thus negatively correlated with Hollingshead score. Within the cross-sectional sample of 623 individuals, SES and IQ were significantly, positively correlated, *r*(621) = 0.31, *p* < 0.001.

### SES and Measures of Brain Volume

We found a strong positive association between SES and total brain volume (*β*_1A_ = 1217.0, *p* < 0.001). A positive association with SES was also seen for total white and grey matter volumes, as well as for total volume of the cortical sheet (Table S3). Separate examination of the two determinants of cortical volume identified a strong positive association between SES and total cortical surface area (*β*_1A_ = 144.1, *p* < 0.001), and a weaker positive association between SES and mean cortical thickness (*β*_1A_ = 0.00086, *p* = 0.01). Greater SES was also significantly associated with greater bilateral volume of all subcortical volumes examined except the pallidum (Table S3). Analysis of standardized effect sizes for SES associations with regional brain volumes revealed that SES was most strongly related to total brain volume and total surface area, such that a standard deviation increase in SES was associated with a 0.17 SD increase in each of these metrics (Figure 1A). Among subcortical structures, SES had the greatest effect on thalamic volume, such that a standard deviation increase in SES was associated with a 0.15 SD increase in thalamic volume (Figure 1A).

**Figure 1.**
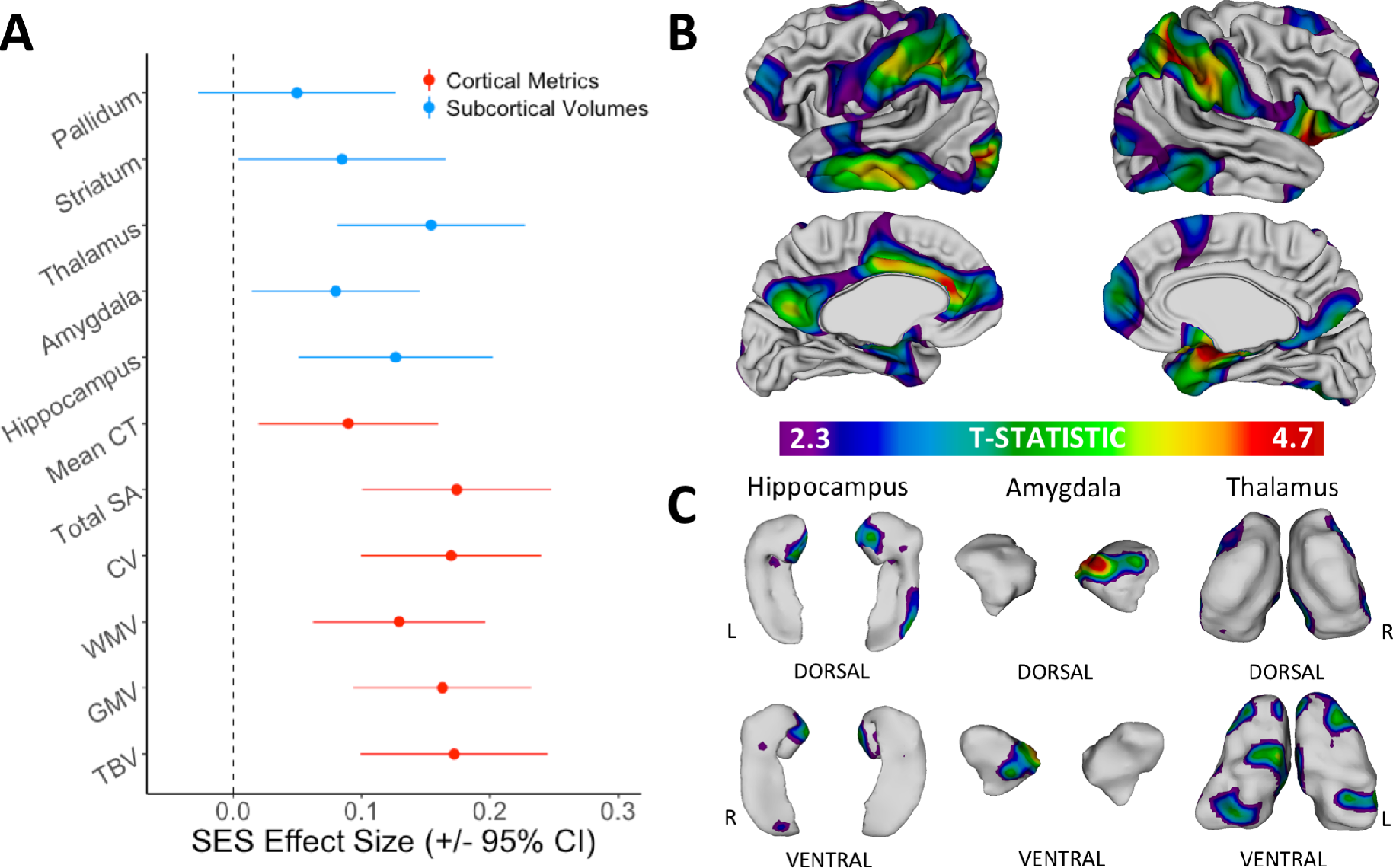
Main effects of SES on global and local anatomy, after controlling for age and sex. (A) Standardized effect size of SES on each global cortical and subcortical brain measure estimated using scaled variables: total brain volume (TBV); grey matter volume (GMV); white matter volume (WMV); cortical volume (CV); total cortical surface area (SA); mean cortical thickness (CT); hippocampus volume; amygdala volume; thalamus volume; striatum volume; and pallidum volume. (B) Cortical surface regions that show a significant positive association with childhood SES. (C) Subcortical surface regions that show a significant positive association with childhood SES.

These above associations between SES and brain anatomy were all developmentally fixed rather than developmentally-dynamic in nature, and comparable between males and females, with two exceptions: (i) the association between SES and total cortical surface area was modified by sex, such that there was a stronger SES effect on surface area for males than females (SES × sex interaction effect: *β*_interaction_ = 122.2, *p* = 0.019; SES main effects: *β*_1A,MALES_ = 198.7, *p* < 0.001/*β*_1A,FEMALES_ = 77.6, *p* = 0.066), and (ii) the association between SES and hippocampal volume was modified by age such that the effect of SES on hippocampal volume grows with age (SES × age interaction effect: *β*_interaction_ = 0.19, *p* = 0.003).

### SES and Cortical Morphology

After establishing that SES was associated with total surface area and mean cortical thickness (Table S3), we tested for regional specificity of these associations through vertex-level analysis of SES associations at 80,962 points (vertices) across the cortical sheet. Vertex-level analyses established that the robust association between greater SES and greater total cortical SA was underpinned by statistically-significant positive associations between SES and regional SA within a distributed set of largely bilateral cortical areas including the lateral prefrontal, anterior cingulate, lateral temporal and superior parietal lobule regions (Figure 1B, Table 2). Associations between SES and vertex-level SA were not significantly modified by age or sex after correction for multiple comparisons (Figure S1).

**Table 2.** Locations of peak SES associations with cortical morphology and top functional associations as indicated by Neurosynth.

| Region of peak vertex | *t* | MNI<br>coordinates |  |  | Top Neurosynth terms |
| --- | --- | --- | --- | --- | --- |
|  |  | x | y | z |  |
| Surface Area |  |  |  |  |  |
| R postcentral gyrus | 4.7 | 27 | -35 | 72 | Motor; Movement; Somatosensory; Execution; Motor Imagery |
| R orbitofrontal cortex | 4.6 | 20 | 19 | -22 | Emotion; Reward; Affective; Fear; Regulation |
| L occipital pole | 4.6 | -35 | -85 | 7 | Visual; Objects; Motion; Orthographic; Reading |
| L inferior parietal cortex | 4.6 | -44 | -38 | 49 | Execution; Motor; Movement; Working Mem.; Spatial |
| L anterior cingulate gyrus | 4.4 | -4 | 36 | 10 | Default Mode (DM); Reward; Self Referential; Emotional; Autobiographical |
| L middle temporal gyrus | 4.3 | -63 | -43 | -17 | Semantic; Retrieval; Word; Mem.; Sentence |
| L precuneus | 3.9 | -4 | -59 | 29 | DM; Theory of Mind; Self Referential; Autobiographical Mem.; Mem. Retrieval |
| R middle temporal gyrus | 3.5 | 61 | -42 | -10 | Semantic; Language; Comprehension; Sentence; Word |
| R precuneus | 3.4 | 4 | -68 | 28 | DM; Autobiographical Mem.; Mem. Retrieval; Self Referential; Episodic Mem. |
| R superior frontal gyrus | 3.4 | 5 | 62 | -5 | DM; Autobiographical Mem.; Self Referential; Social; Theory of Mind |
| Cortical Thickness |  |  |  |  |  |
| R supramarginal gyrus | 4.3 | 59 | -19 | 33 | Somatosensory; Motor; Tactile; Movement; Pain |

Associations between SES and regional cortical thickness were much more localized than observed for regional SA. Specifically, we identified a single locus of significant positive association between SES and CT in the right supramarginal gyrus (Figure S2, Table 2).

### SES and Subcortical Morphology

We investigated the spatial specificity of observed associations between SES and bilateral subcortical volumes by modeling surface area at each vertex across the surface of the hippocampus, amygdala, thalamus, and striatum (Figure 1C). Hippocampal effects were concentrated bilaterally on the medial surface, and neighboring medial surfaces of the right amygdala also showed a significant positive association with SES. The association between SES and thalamic surface area was localized primarily to the ventral posterior and ventral lateral thalamus. No other subcortical structures showed statistically significant shape associations with SES after correcting for multiple comparisons across vertices.

### Separating Main Effects of SES and IQ on Anatomy

Because of the noted strong association between SES and IQ (Figure 2A), we next re-ran linear mixed-effects models with Full-Scale IQ as a covariate, in order to parse main effects of SES and cognition on neuroanatomy. We observed positive associations between IQ and all global measures of cortical and subcortical anatomy (Table S3). The main effect of SES on each global metric was reduced in magnitude by the addition of IQ to the model; for mean cortical thickness, bulk amygdalar volume, and bulk striatal volume, the main effect of SES was no longer significant (Table S3), although SES continued to show a significant positive association with TBV, GMV, WMV, CV, total cortical SA, and bilateral hippocampal and thalamic volumes after controlling for IQ. There were no significant SES × IQ interactive effects on any global metrics.

**Figure 2.**
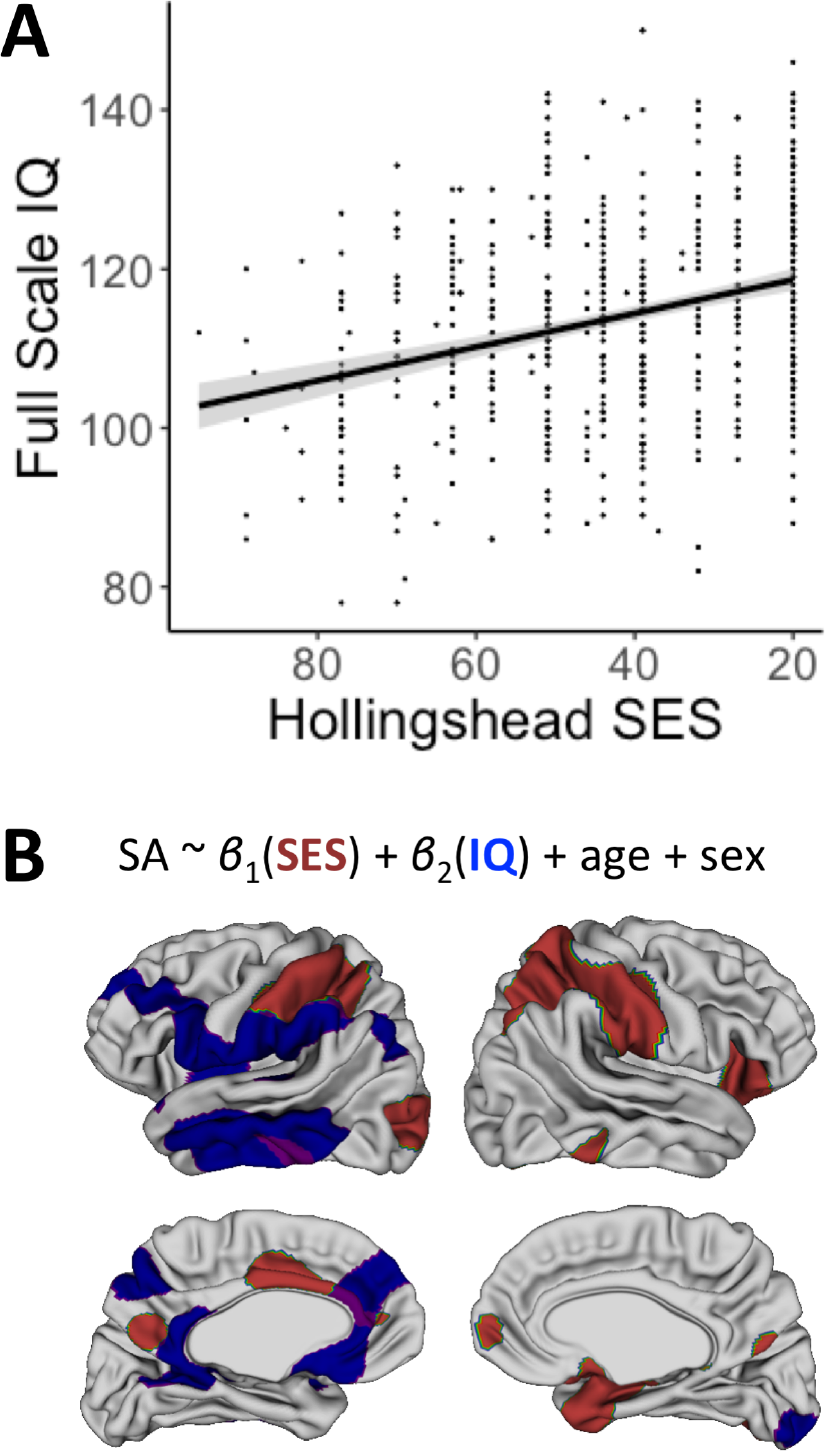
(A) Correlation between childhood SES and Full-Scale IQ in cross-sectional sample of 623 individuals, *r*(621) = 0.31, *p* < 0.001. (B) Map of main effects of SES and IQ on cortical surface area. Significant SES main effect is represented in red, IQ main effect in blue, and overlapping main effects in purple. Together, these main effects resemble the map of main SES effects on cortical SA without IQ as a covariate (Figure 1B).

We next probed the spatial patterning of IQ versus SES main effects on cortical surface area. After false discovery rate (FDR) correction for multiple comparisons, childhood SES and IQ were both positively associated with regional cortical SA (Figure 2B). Specifically, when including IQ as a covariate, the main effect of SES was restricted to the bilateral superior parietal, right orbitofrontal, left inferior temporal, and bilateral medial prefrontal cortices. The main effect of IQ, in contrast, was localized to the left inferior and middle temporal, left inferior parietal, and left medial frontal regions. Strikingly, the regional maps of separate SES and IQ effects on cortical surface area together (Figure 2B) resemble the regional effects of SES when IQ is not included as a covariate (Figure 1B). In other words, the main effect of SES on cortical surface area appears to be separable into effects related to the strong IQ differences across SES, localized to the left-lateralized perisylvian and medial prefrontal cortices, and SES-SA associations that are independent from associated variation in general cognitive ability, localized to bilateral parietal and frontal cortices.

### Structural Measures Mediate the Association Between SES and IQ

In addition to testing the effects of IQ as a covariate in SES-brain associations, we also probed the relation between SES, cognition, and anatomy by examining whether any of the structural brain measures found to be significantly associated with SES might mediate the association between SES and IQ in our cross-sectional dataset of 623 individuals. Using the Mediate package in R (Tingley et al. 2014), we estimated the proportion of the total relation between SES and IQ that was accounted for by each anatomical mediator. Among global cortical measures, total brain volume (proportion mediated = 0.148, *p* < 0.001), grey matter volume (proportion mediated = 0.145, *p* < 0.001), white matter volume (proportion mediated = 0.077, *p* < 0.001), cortical volume (proportion mediated = 0.146, *p* < 0.001), total surface area (proportion mediated = 0.114, *p* < 0.001), and mean cortical thickness (proportion mediated = 0.049, *p* = 0.01), each partially mediated the relation between SES and IQ. Subcortical volumes also partially mediated the association (hippocampus, proportion mediated = 0.052, *p* < 0.001; amygdala, proportion mediated = 0.034; *p* = 0.02; thalamus, proportion mediated = 0.087; *p* < 0.001; striatum, proportion mediated = 0.033, *p* = 0.01).

Finally, we extended the mediation analysis to investigate whether there was a regional specificity to the partial mediation of SES and IQ by cortical and subcortical surface area. To do so, we submitted all of the vertices in each hemisphere that showed a significant association with SES to a multiple mediation analysis using the MultiMed package in R (Boca and Sampson 2014). Three cortical regions – all in the left hemisphere – showed significant mediation of the SES association with IQ: the middle temporal gyrus, supramarginal gyrus, and anterior cingulate cortex (Figure S3). Notably, these largely overlapped with the cortical vertices that showed shared main effects of SES and IQ on surface area (regions in purple, Figure 2B).

### Functional Implications of Anatomical Results

Coordinates of peak regional SES associations with cortical surface area and cortical thickness were submitted to Neurosynth (Yarkoni et al. 2011), a meta-analytic repository of structure-function relationships. The top five cognitive and psychological terms with the highest meta-analytic co-activation correlation coefficient *r* are displayed in Table 2, after redundant terms and anatomical terms were removed. This platform provided evidence that the spatial effects of SES are localized to regions preferentially associated with sensorimotor functions, as well as language, memory, and emotional processing, which have each been shown to exhibit SES differences (Farah et al. 2006; Noble et al. 2007; Kim, Evans, et al. 2013).

## Discussion

Here, with a large, longitudinal, single-site neuroimaging sample, we both replicate and extend findings on the relation between childhood SES and structural brain anatomy in a number of key directions.

First, in line with previously replicated findings, we find that higher SES is associated with greater global brain measures including greater grey matter volume (Hanson et al. 2013; Luby et al. 2013; Mackey et al. 2015; Gianaros et al. 2017), greater cortical surface area (Noble et al. 2015; Gianaros et al. 2017), greater cortical thickness (Mackey et al. 2015; Noble et al. 2015) and greater hippocampal volume (Noble et al. 2012, 2015; Luby et al. 2013; Hair et al. 2015). While previous findings regarding white matter volume and amygdala volume have been mixed, with some studies reporting positive associations with SES and others reporting no significant association (Jednoróg et al. 2012; Luby et al. 2013; Noble et al. 2015; Gianaros et al. 2017), we find a significant positive SES association with both white matter volume and amygdala volume. We further extend analyses of SES associations with subcortical anatomy to provide the first evidence that greater childhood SES is associated with larger bilateral volumes of the thalamus and striatum. The thalamic finding is especially notable given reported associations between thalamic volume and cognitive measures including cognitive speed and verbal IQ (Van Der Werf et al. 2001; Xie et al. 2012). Comparative analyses of SES associations using standardized effect sizes ranks total cortical surface area and bilateral thalamic volume as the morphometric properties of the cortex and subcortex, respectively, that show the largest effect-size relations with SES.

Second, our study advances understanding of the neuroanatomical correlates of SES by using surface-based algorithms to further parse associations between SES and anatomy of the cortical sheet. We observe SES associations with cortical surface area in a number of largely bilateral regions including lateral prefrontal, anterior cingulate, lateral temporal, and superior parietal lobule regions, as well as with cortical thickness in the right supramarginal gyrus. These SES-surface area associations largely correspond to the map of associations between parental education and cortical surface area presented by Noble and colleagues (Noble et al. 2015). Functional interpretation with Neurosynth (Yarkoni et al. 2011) suggests that the cortical regions we find to show morphological associations with SES are preferentially involved in networks that underlie sensorimotor functions, as well as language, memory, and emotional regulation.

Third, we provide the first systematic examination of SES associations with subcortical shape, which indicates that variations in childhood SES are associated with focal differences in hippocampal, amygdalar and thalamic anatomy. Lower SES was associated with reduced surface area in medial amygdalo-hippocampal sub-regions, and associations between SES and thalamic shape localized to regions overlying the ventral lateral and ventral posterior nuclei (Figure 1C). Histological studies and functional connectivity neuroimaging studies suggest that these thalamic nuclei are preferentially connected to frontoparietal cortical systems which subserve primarily sensorimotor functions (Jones 1985; Kim, Park, et al. 2013). Strikingly, we also detect strong associations between SES and surface area within these cortical targets of ventral thalamic nuclei, suggesting that anatomical correlations of childhood SES variation may be organized by the topography of cortico-subcortical connectivity.

Finally, we demonstrate convergent anatomical correlates of the strong relation between SES and IQ using two independent methods. Modeling the main effects of both IQ and SES on vertex-wise surface area revealed that the map of statistical SES effects can be fractionated into distinct SES and IQ effects. Additionally, we demonstrate that many of the anatomical correlates of SES significantly mediate the relation between SES and IQ. We expand upon previous results that found significant SES-cognition mediation by whole-brain surface area (Noble et al. 2015) by conducting cross-sectional mediation tests at each cortical vertex that had a significant SES-surface area association. Specifically, we find that vertices in the left middle temporal gyrus, left supramarginal gyrus, and left anterior cingulate – the same three regions that showed a main effect of IQ when controlling for SES – mediate the relation between SES and IQ. These three regions are known to be involved in language development (Vannest et al. 2009) and cognitive control (Shackman et al. 2011).

A mediation analysis allows us to test one possible pathway between SES, brain anatomy, and cognition, and we show that SES may exert some of its effect on cognition by altering structural brain development, particularly in regions associated with language and learning. However, it is important to note that this pathway represents only one possible set of interactions between childhood environment, anatomy, and cognition. Farah (Farah 2017) provides a succinct review of the main processes that may operate to exacerbate neural and psychological SES disparities: the social causation hypothesis suggests that the environmental conditions associated with different levels of SES influence brain structure and function, while the social selection hypothesis suggests that genetic factors in parents that both affect their cognition and predispose them to a certain SES level are then transmitted to their children. These processes likely both operate in concert and interact to some degree. Because of the inherent observational nature of studies of SES, it is difficult to determine to what extent the anatomical correlates of SES that we report here may reflect the shared genetic effects on SES and brain development versus direct effects of SES on brain development. Nevertheless, our findings help to pinpoint cortical and subcortical systems which represent candidate biological substrates for these diverse causal pathways.

### Caveats and Future Directions

One important limitation of this sample is the composite nature of the Hollingshead 2-factor index of SES. The field has lately recognized the importance of differentiating between the effects of separate components of socioeconomic status (Duncan and Magnuson 2012). A few studies have already begun to identify the differential effect that parental education and family income may have on structural brain development (Lawson et al. 2013; Noble et al. 2015). Unfortunately, when data collection for this sample began in the 1990s, the Hollingshead was selected for its widespread use and ease of measurement, and only the single, derived SES score was recorded initially. As future studies are designed it will be important to report on more nuanced factors that compose SES and may have unique effects on brain development and thus serve as specific targets for intervention.

It is also important to note that the subjects in this sample are not representative of the socioeconomic distribution in the United States. Sample composition is known to affect conclusions about the normative trajectory of brain development (LeWinn et al. 2017). Additionally, some studies have suggested that socioeconomic variables relate most strongly to brain development among the most disadvantaged children (Noble et al. 2015), and the fact that we did not detect such a gradient may be due to our unrepresentative SES distribution. The average IQ of our subjects was also higher than the expected mean of 100 (Table 1); however, IQ is known to increase across generations in the general population, so the high mean IQ may be partially attributable to the use of the same IQ test across the multiple decades of this study (Flynn, 1987). Although we cannot generalize across the socioeconomic or cognitive spectrums, it is notable that we found that SES variation is related to detectible differences in cognitive ability and structural brain development within a typically developing cohort lacking frank socioeconomic deprivation.

### Conclusion

Childhood socioeconomic status is a complex construct that influences the physical and psychosocial environment in which a child develops. Here, we demonstrate regionally-specific associations between childhood SES and both cortical and subcortical morphology. Our findings inform ongoing efforts to clarify the spatiotemporal patterning of SES-related neuroanatomical variation and its relation to cognitive outcomes such as IQ. Of note, the results presented here do not establish a direct causal pathway between SES and brain development, nor do they indicate that childhood SES exerts a deterministic effect on development. Rather, by resolving neuroanatomical substrates that vary closely with SES, we contribute new biological information to a growing field of multidisciplinary research that ultimately aims to reduce SES variation in health and achievement.

## Acknowledgements

The authors wish to thank the participants and families who took part in this study. The authors have no conflicts of interest to declare.

